# Whole genome duplication in coast redwood (*Sequoia sempervirens*) and its implications for explaining the rarity of polyploidy in conifers

**DOI:** 10.1101/030585

**Authors:** Alison Dawn Scott, Noah Stenz, David A. Baum

## Abstract

- Whereas polyploidy is common and an important evolutionary factor in most land plant lineages it is a real rarity in gymnosperms. Coast redwood (*Sequoia sempervirens*) is the only hexaploid conifer and one of just two naturally polyploid conifer species. Numerous hypotheses about the mechanism of polyploidy in *Sequoia* and parental genome donors have been proffered over the years, primarily based on morphological and cytological data, but it remains unclear how *Sequoia* became polyploid and why this lineage overcame an apparent gymnosperm barrier to whole-genome duplication (WGD).
- We sequenced transcriptomes and used phylogenetic inference, Bayesian concordance analysis, and paralog age distributions to resolve relationships among gene copies in hexaploid coast redwood and its close relatives.
- Our data show that hexaploidy in the coast redwood lineage is best explained by autopolyploidy or, if there was allopolyploidy, this was restricted to within the Californian redwood clade. We found that duplicate genes have more similar sequences than would be expected given evidence from fossil guard cell size which suggest that polyploidy dates to the Eocene.
- Conflict between molecular and fossil estimates of WGD can be explained if diploidization occurred very slowly following whole genome duplication. We extrapolate from this to suggest that the rarity of polyploidy in conifers may be due to slow rates of diploidization in this clade.

## Introduction

Polyploidy has profound long- and short-term genetic consequences (Adams & Wendel, 2005; Otto & Whitton, 2000; etc.), and facilitates adaptive evolution (Soltis et al., 2008; etc). Studies of genome sequences, expressed genes, and cytogenetics suggest that all land plant lineages have experienced polyploidization in their evolutionary history, though clades differ in the extent of recent whole genome duplication (neopolyploidization). While there are thousands of neopolyploid mosses, ferns and angiosperms, the phenomenon is relatively rare in gymnosperms, and especially conifers. There are only two polyploid conifer species: alerce, *Fitzroya cupressoides* (4x), and coast redwood, *Sequoia sempervirens* (6x). Why is polyploidy so rare in conifers? Does it reflect rare formation of polyploid individuals, for example due to a lack of unreduced gametes, or another barrier to allopolyploid formation? Or, do polyploid taxa form in gymnosperms, but fail to give rise to successful clades? To shed light on these questions, we studied the evolutionary history of coast redwood with the goal of determining when polyploidy occurred and whether it entailed allopolyploidy.

Coast redwoods are long-lived trees (some over 2,000 years; Burns & Honkala, 1990) that thrive in the foggy coastal forests of central and northern California. Coast redwoods are among the world’s tallest living trees (up to 115 meters; Ishii et al., 2014). *Sequoia* is a monotypic genus whose closest relatives are the giant sequoia of the Californian Sierra Nevada (*Sequoiadendron giganteum*) and the Chinese dawn redwood (*Metasequoia glyptostroboides*). Though the three modern redwood species have distinct ranges, fossil data suggest that diverse redwood lineages were widely distributed across the Northern Hemisphere from the Cretaceous onwards (Miller, 1977). The oldest redwood fossils are from South Manchuria (present-day China) and Boulogne-sur-Mer (northern France) and date back to the mid-to-late Jurassic, suggesting the redwood clade is at least 146 million years old (Zeiller and Fliche, 1903; Endo, 1951).

*Sequoidendron* and *Metasequoia* are diploids with 2n=22 (Schlarbaum and Tshuchiya, 1984). Hirayoshi and Nakamura (1943) first determined the correct chromosome number of *Sequoia* and proved that it is a hexaploid with 2n=66. Hexaploidy in *Sequoia* was later corroborated by Stebbins (1948), Saylor and Simons (1970) and Ahuja and Neale (2002).

Relying on the well-known correlation between guard cell size and genome size (e.g., Beaulieu et al., 2008), Miki and Hikita (1951) studied stomatal guard-cell size in Pliocene fossils of Metasequoia and Sequoia. As fossil guard cells were the same size as extant guard cells, Miki and Hikita concluded *Sequoia* has been hexaploid since at least the Pliocene (2.5-5 million years ago). This estimate was pushed back significantly by Ma et al. (2005), who describe fossils from the Eocene (33-53mya) with guard cells of a size taken to indicate polyploidy.

Morphological similarities among modern redwoods led to hypotheses of allopolyploidy in *Sequoia* involving hybridization between extinct diploid *Sequoia* and ancestors of either *Metasequoia* (Stebbins, 1948) or *Sequoiadendron* (Doyle, 1945). Despite the distance among their modern ranges, the overlap in fossil distributions of *Sequoia, Sequoiadendron,* and *Metasequoia* make this hypothesis plausible. Another hypothesis is that an extinct member of the Taxodiaceae, perhaps a member of *Taxodium,* contributed to the hexaploid genome of *Sequoia* (Stebbins, 1948; Saylor and Simons, 1970). Ahuja and Neale (2002), in contrast, suggested that the “missing” parent of *Sequoia* may have been a member of the *Cryptomeria, Taiwania,* or *Athrotaxis* lineages.

Before the advent of molecular phylogenetics, auto- and allopolyploids were distinguished by observing chromosome behavior during meiosis. Autopolyploidy (generally interpreted as occurring within a single species) and allopolyploidy (involving hybridization among species) represent extremes of a spectrum. Autopolyploids have multiple sets of very similar homologous chromosomes, which tends to manifested cytogenetically as the formation of multivalents (e.g. groups of four or six chromosomes). Allopolyploids, in contrast, arise from the fusion of divergent genomes which, in the extreme case results in bivalent formation by each homologous chromosome, as observed in diploid organisms. However, chromosome pairing at meiosis is rarely definitive as allopolyploidy can result in multivalent formation among homeologs if hybridizing species are closely related, and bivalent formation is eventually reestablished following autopolyploidy by the process of diploidization (Ramsey and Schemske, 2002; Parisod et al., 2010).

In addition to cytogenetic lines of evidence, segregation patterns can be useful to distinguish auto- and allopolyploids. An autopolyploid forming multivalents at meiosis will produce equal frequencies of all possible allele combinations. In the case of *Sequoia,* this pattern is called hexasomic inheritance. Allopolyploids do not typically form multivalents at meiosis, resulting in simple disomic inheritance (as seen in diploids). Again, these are only the most extreme possibilities, as both the diploidization process and polyploidy involving a mixture of similar and divergent chromosomes (i.e. segmental allopolyploidy sensu Stebbins) can lead to intermediate inheritance patterns.

Studies of meiotic chromosome pairing in *S. sempervirens* reported a mixture of bivalents and multivalents (Stebbins, 1948; Schlarbaum and Tsuchiya, 1984; Ahuja and Neale, 2002). This led Stebbins (1948) and Schlarbaum and Tsuchiya (1984a, b) to suggest that hexaploidy involved both auto- and allopolyploidy. A similar result was obtained by Rogers (1997), who used allozymes to study inheritance patterns in *Sequoia.* However, neither the pairing nor genetic data are sufficient to distinguish segmental allopolyploidy from autoployploidy followed by partial diploidization We set out to use modern genomic approaches to revisit the evolutionary history of polyploidy in *S. sempervirens* and see if, by doing so, we could also gain insights into why polyploidy is so rare in gymnosperms.

## Materials and Methods

### Transcriptome sequencing and assembly

Total RNA was extracted from foliage samples of *S. sempervirens, S. giganteum, M. glyptostroboides,* and the outgroup *Thuja occidentalis* (eastern white cedar) with a CTAB/Chisam extraction protocol followed by Qiagen RNeasy cleanup. Illumina TruSeq cDNA libraries were prepared and sequenced on an Illumina HiSeq 2000 with 100bp paired-end reads at either the UW Biotech Center (Madison, WI) or at the SciLife Laboratory (Stockholm, Sweden).

### Sequence analysis and alignment

We assembled raw reads *de novo* with Trinity vers. 2014-07-17 (Grabherr et al., 2011), with default settings and Trimmomatic processing. After assembly, contigs were translated using TransDecoder vers. 2014-07-04 (Haas et al., 2013; http://transdecoder.sf.net) with a minimum protein length of 100aa. Translated contigs were filtered using the Evigene pipeline vers. 2013.07.27 (http://arthropods.eugenes.org/EvidentialGene/about/EvidentialGene_trassem_blypipe.html). Ortholog clusters shared among *S. sempervirens, S. giganteum, M. glyptostroboides,* and *T. occidentalis* were identified using the translated transcriptome assemblies by ProteinOrtho ver. 5.11 (Lechner et al., 2011), using an algebraic connectivity cutoff of 0.25. Custom Perl scripts (available at github.com/nstenz) were used to identify ortholog sets that contained a single copy in diploids (*S. giganteum, M. glyptostroboides,* and *T. occidentalis)* and between one and three copies in the hexaploid *S. sempervirens*. As these putatively single-copy protein-coding sequences show marked conservation among species, we assumed that allelic variants would generally be combined into a single contig. We used MUSCLE v. 3.8.13, 64bit (Edgar, 2004a,b), with default alignment settings to align the ortholog sets at the protein level before using a custom PERL script to generate the corresponding nucleotide alignment.

### Single-variant gene trees and concordance analyses

For each orthogroup that included only one sequence variant in *S. sempervirens* we estimated phylogenetic trees using MrBayes vers. 3.2.2 64bit (Huelsenbeck & Ronquist, 2001; Ronquist & Huelsenbeck, 2003) with the settings: nst = 6; rates = invgamma; ngen = 1.1 million; burnin = 100,000; samplefreq = 40; nruns = 4; nchains = 3; temp = 0.45; swapfreq = 10. BUCKy vers. 1.4.4 (Ané et al., 2007; Larget et al., 2010) was then used to estimate the proportion of genes that have each possible resolution in the redwood clade while taking account of uncertainty in individual gene trees. Post-burnin posterior distributions from MrBayes were combined in BUCKy for 1 million generations with α = 1. All trees were rooted on the outgroup, *Thuja occidentalis.*

### Density distribution of K_s_ estimates

To build an age distribution of K_s_ (the average number of synonymous substitutions per synonymous site) within each transcriptome we identified duplicate genes using custom Perl scripts (available at github.com/nstenz). Assembled contigs were translated using TransDecoder with a minimum protein length of 100aa, as above. Duplicate genes were identified using BLAT (Kent 2002) on translated contigs and then duplicate gene pairs were aligned and back translated into their corresponding nucleotide sequence. We estimated K_s_ on each pair of nucleotide alignments using K_a_K_s_ calculator (model GY; Zhang et al., 2006). We excluded K_s_ values greater than 2 to avoid the effects of K_s_ saturation, and plotted the resulting K_s_ values in a density plot in R (R core team, 2013). To identify significant features of the K_s_ frequency distributions we used SiZer (Chaudhuri and Marron, 1999).

### Multi-variant gene trees and tree-based K_s_ estimates

For alignments containing a single variant in diploid taxa and two or three variants in hexaploid *Sequoia,* we estimated phylogenetic trees with raxml vers. 8.1.20 (100 bootstrap replicates; GTRGAMMA; Stamatakis, 2006). We then used PAML (Yang, 1997) to obtain a tree-based estimate of K_s_. PAML calculates branch lengths along the ML tree using a model that estimates the rate of synonymous and non-synonymous substitutions (D_s_ and D_n_, respectively) separately for each branch. We imposed a molecular clock assumption (clock=1) to obtain an ultrametric tree. By multiplying a branch’s length by its D_s_ and summing over intervening branches between two tips we could obtain an estimate of the patristic K_s_ distance between *Sequoia* homeologs and how this compares to the K_s_ of copies from different species.

In order to obtain an approximate date for gene duplication, we divided the depth of the gene duplication in K_s_ units by an average mutation rate for conifers of 0.68×10^−9^ synonymous substitutions per synonymous site per year (Buschiazzo et al., 2012). *Sequoia* is hexaploid, so at least two whole genome duplications must have occurred in the past. As each whole genome duplication event is expected to yield a normal distribution of K_s_ values, we used EMMIX v.1.3 (Mclachlan et al., 1999) to fit a mixture model of normal distributions as a way to assign putative homeologs to each duplication event and estimate their ages. We allowed EMMIX to fit 1-2 normal distributions, with the optimal model selected based on AIC and BIC scores.

## Results

Our *de novo* transcriptome assemblies ranged from 70 to 101mbp in length (Table 1). Assembled contigs per species ranged from 80,126 to 128,005.

**Table 1.** Assembly statistics

| Taxon | Raw reads<br>(paired end) | Assembly length<br>(mbp) | Contigs | N50 |
| --- | --- | --- | --- | --- |
| *Sequoia sempervirens* | 55,052,935 | 85.6 | 128,005 | 1,118 |
| *Sequoiadendron giganteum* | 56,665,524 | 101.3 | 115,519 | 1,619 |
| *Metasequoia glyptostroboides* | 29,502,075 | 78.6 | 83,120 | 1,668 |
| *Thuja occidentalis* | 31,116,702 | 70.0 | 80,126 | 1,607 |

Assuming synonymous substitutions happen at a constant rate over time, Ks can be used as a proxy for the age of duplicate genes. To estimate the distribution of pairwise K_s_ distance within each genome, we identified all duplicate genes, which numbered 33,544, 39,236, and 26,485, in *S. sempervirens, S. giganteum,* and *M. glyptostroboides,* respectively. Paralog age distribution plots for all three taxa revealed a peak at a K_s_ ~ 1.5, of which those for *S. sempervirens, S. giganteum* are shown in Fig. 1. Allowing for the approximate nature of these calculations, this peak likely corresponds to the seed plant whole genome duplication previously dated at 319 Ma (Jiao et al., 2011). Despite the expectation that hexaploid *Sequoia* would have at least one other, much younger peak corresponding to a polyploidization event in perhaps the Eocene (Ma et al., 2005), this was not visible in the age distribution plots (Fig. 1). Results from SiZer also did not indicate any significant peak unique to the *Sequoia* K_s_ plot.

**Figure 1:**
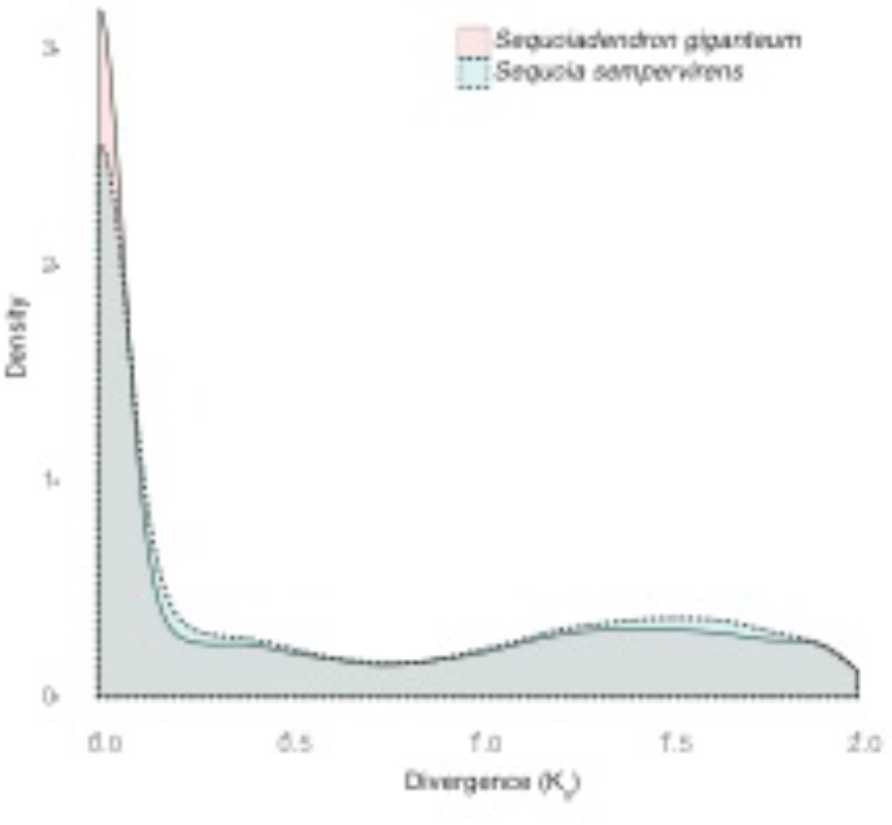
Density distribution of pairwise Ks between duplicate genes in *Sequoia* (pink) and *Sequoiadendron* (cyan).

To distinguish the evolutionary relationships among redwoods and look for evidence of ancestral hybridization, we used Bayesian concordance analysis and estimated genomic support for each of three possible topologies for an unrooted four-taxon tree. First we built individual gene trees from 7,819 ortholog groups that each had one sequence variant in each diploid species (*Sequioadendron, Metasequoia, Thuja*) and one, two, or three sequence variants in the hexaploid, *Sequoia.* Alignment lengths in this set varied from 301-5,736 bp, with a median of 1,104. Of these alignments 7,602 included a single *Sequoia* copy, whereas 217 included one or two *Sequoia* sequence variants. Among the 7,602 alignments that included a single copy in *S. sempervirens* the most frequently supported topology placed *S. sempervirens* sister to *Sequoiadendron* (Fig. 2) with a concordance factor (CF; Baum 2007) mean estimate of 0.79 and a 95% credibility interval of 0.78-0.80. The two minor topologies (*Sequoia* + *Metasequoia*; *Metasequoia* + *Sequoiadendron*) had concordance factors of 0.10(0.09-0.11) and 0.11(0.10, 0.12), respectively (Fig. 2). These results show that, if *Sequoia* arose from allopolyploidy, it only involved genome donors in the Californian redwood clade (i.e., the clade that includes *S. sempervirens* and *Sequoiadendron*). However, autoploidy is also a possibility.

**Figure 2:**
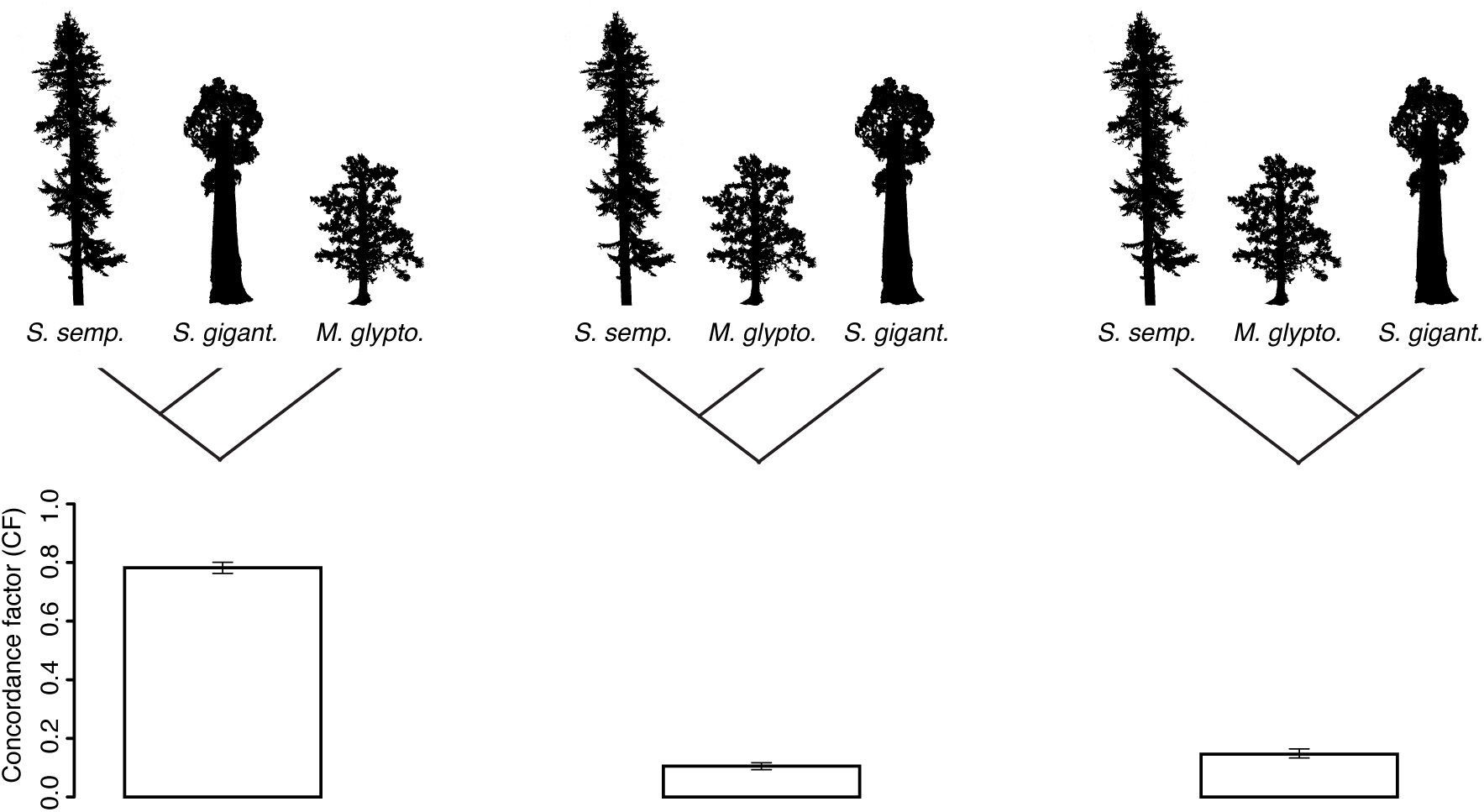
Bayesian concordance analysis of 7,602 gene trees. For each of three possible topologies, the concordance factor (proportion of loci in the sample having the clade) and its 95% credibility interval are shown.

In order to obtain estimates for the divergence of *Sequoia* duplicates relative to interspecies divergences and to re-evaluate evidence for allopolyploidy within the Californian redwood clade, we estimated phylogenetic trees for all genes with more than one sequence variant in *Sequoia.* A total of 217 genes were present in two or three copies in *S. sempervirens.* The optimal tree for 186 of these alignments (85.7%) showed monophyly of the *S. sempervirens* copies with *Sequoia* sister to *Sequoiadendron* (Fig. 3), with 97% of these trees well-supported (i.e., having a bootstrap > 0.70). The remaining 31 genes (14%) either contradicted monophyly of *S. sempervirens* copies, supporting several other possible relationships, or lacked clear resolution of species relationships.

**Figure 3:**
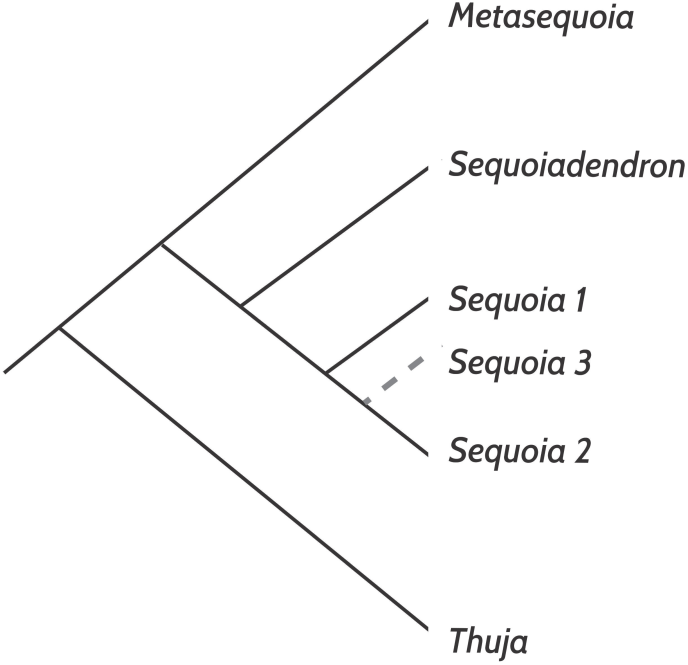
Cladogram summarizing 184 gene trees as estimated by MrBayes.

Based on ML estimates using a codon model in PAML, we could calculate the patristic Ka and Ks distances between each pair of tips for on each genes tree. Doing this on the 176 well-supported gene trees that yielded a monophyletic *Sequoia,* average phylogenetic K_s_ among *Sequoia* gene copies was 0.013. This was approximately one-third of the Ks separating *Sequoia* sequences from other redwoods (Figure 4).

**Figure 4:**
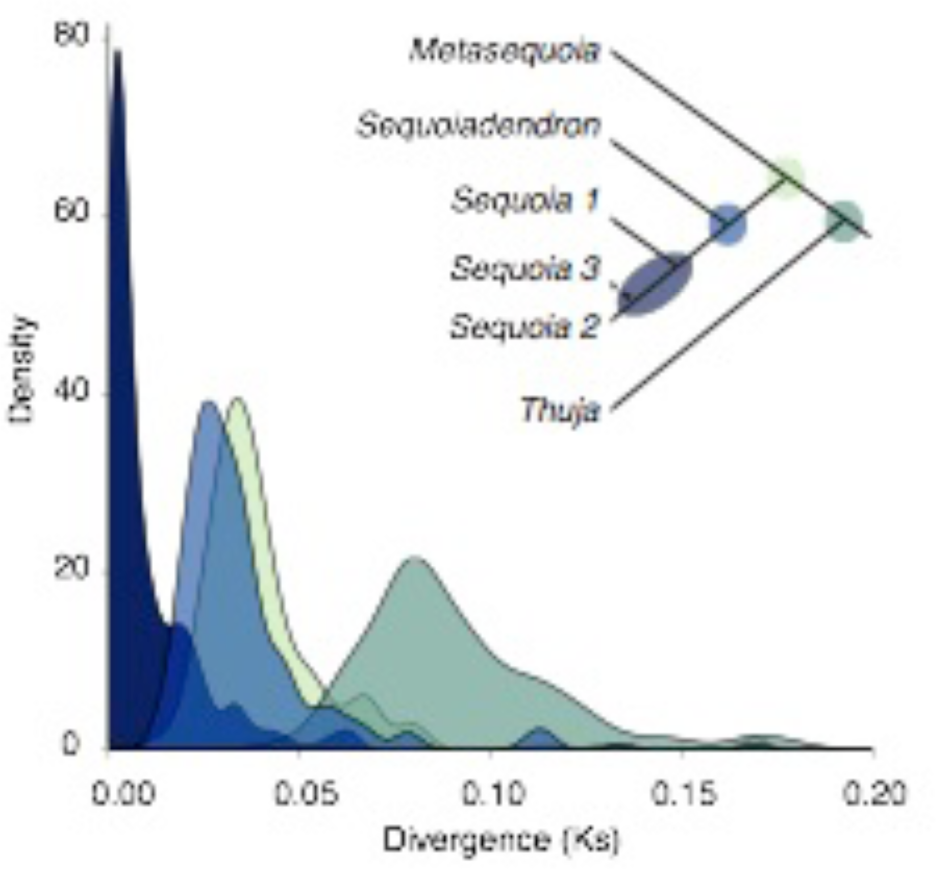
Tree-based divergence estimates in Ks. Density distribution of divergence estimates (in Ks). For Distributions are colored to indicate corresponding nodes on the tree.

We tested whether the patristic Ks estimates between *S. sempervirens* copies are sampled from one or two normal distributions. If hexaploidy arose from two sequential WGD events, there should be two, distinct normal distributions. We used EMMIX to fit a mixture model of normal distributions to the PAML Ks estimates. Based on AIC and BIC scores, the presence of two Gaussian distributions provides a better fit to the K_s_ distance data. Figure 5 shows the best fitting pair of distributions. Although it is difficult to reliably translate Ks into absolute age, using a generic average mutation rate for conifers of 0.68×10^−9^ synonymous substitutions per site per year (Buschiazzo et al., 2012), these peaks correspond to ~3 Ma and 10 Ma.

**Figure 5:**
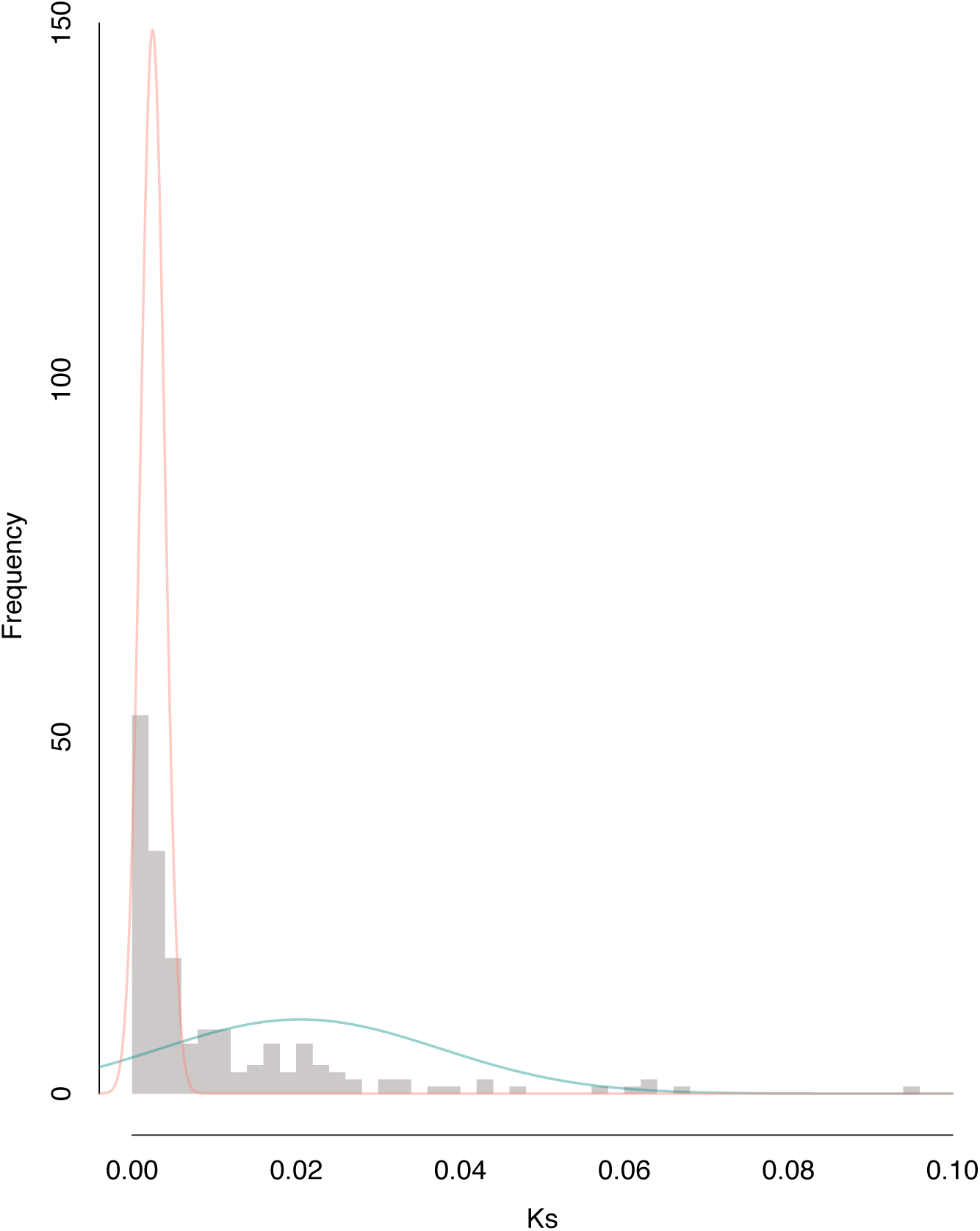
Age distribution of *Sequoia* variants. Colored lines denote normal distributions fit with EMMIX.

## Discussion

### Transcriptome sequencing in the redwoods supports a sister group relationship between *Sequoia* and *Sequoiadendron.*

Bayesian concordance analysis of single copy genes overwhelmingly supports *Sequoiadendron* as the closest relative of *Sequoia.* This conclusion is in agreement with decades of previous work based on morphology, karyotype, and chloroplast sequence data (e.g. Brunsfield et al., 1994; Gadek et al., 2000; Kusumi et al., 2001).

We found genes supporting two minor topologies, one with a *Sequoia-Metasequoia* clade and the other with a *Sequoiadendron–Metasequoia* clade. These discordant topologies could be due to incomplete lineage sorting (ILS), which arises when multiple gene copies (or alleles) persist between sequential splits in a population tree. In this case, the two minor trees have similar concordance factors, 0.010 and 0.11, and their associated credibility intervals overlap. This pattern is consistent with ILS, which predicts that the alternative minor topologies should have equal CFs (Baum 2007). Furthermore, given a concordance factor of 0.80, coalescent theory would predict that *Sequoia-Sequoiadendron* clade is subtended by a population lineage whose duration was ~1.22 Ne generations, where Ne is the effective population size (Allman et al., 2011; Larget et al. 2011). However, it is also possible that the internal branch is considerably longer and discordance is due to other factors such as mistaken orthology. The fact that the two minor histories have similar concordance factors tends to argue against introgression or hybridization as an important phenomenon in the group.

### Hexaploidy in *Sequoia* did not involve hybridization among extant redwood lineages

Our phylogenetic results support an autopolyploid origin for hexaploid *Sequoia*, with no evidence to support hybridization among modern redwood lineages. Single-copy trees convey strong support for *Sequoiadendron* as the closest relative of *Sequoia,* suggesting there was no genome contribution from *Metasequoia*. The lack of evidence that *Metasequoia* was involved with the polyploid origins of *Sequoia* puts some long-held hypotheses to rest (e.g. Stebbins, 1948; Saylor & Simons, 1970). However, as these phylogenies include only one copy for hexaploid *Sequoia*, they could not distinguish between autopolyploidy within the *Sequoia* lineage or autoallopolyploidy within the *Sequoiadendron-Sequoia* clade. Single-copy trees may also be inconclusive due to extreme copy-specific expression or genome dominance, where genes from one parental genome are preferentially expressed (e.g. Woodhouse et al., 2014). Therefore, we sought additional evidence by studying orthogroups that included 2 or 3 distinct sequence variants, putatively homeologs, from *Sequoia.* Phylogenetic analyses of these orthogroups strongly support monophyly of *Sequoia* homeologs, suggesting that all gene copies in *Sequoia* originate from the same redwood lineage.

### Polyploidy in *Sequoia* arose relatively recently

The similarity of the K_s_ plots obtained from polyploid *Sequoia* and diploids *Sequoiadendron* and *Metasequoia* (Fig. 2), and specifically the lack of a recent peak restricted to *Sequoia*, is initially surprising, as these methods have been widely used to diagnose polyploidization events in numerous plant lineages (e.g. Barker et al., 2008; Jiao et al., 2011). This pattern might be expected if autopolyploidy had occurred very recently, such that the level of divergence among homeologs is not much different than that among alleles at a particular locus (Vanneste et al. 2013), but the fossil data suggests polyploidization as early as the Eocene. One possible explanation for the lack of a polyploidization peak is that only one homeolog is expressed in leaves. Such genome dominance has been observed in other polyploid species (e.g., Adams et al. 2004). However, the fact that we found many genes with two or three distinct copies in *Sequoia* but only one in each diploid argues against uniform silencing of all but one homeolog.

To further explore the history of gene duplication, we inferred trees for alignments that included one transcript in diploids and two or three from *Sequoia* and then inferred the branch lengths of this tree in K_s_ units. We found that K_s_ estimates between even the most divergent *Sequoia* homeologs were very low (>0.10). One possible explanation is that *Sequoia* experienced a long period of multisomic inheritance following autopolyploidy during which time homeologs tended to be repeatedly recombined, resulting in much lower K_s_ values (described in Wolfe, 2001). These observations highlight some caveats of using paralog age distribution graphs alone to infer recent polyploidization events, or to study ancient whole genome duplication events that were accompanied by extended periods of multisomic inheritance.

Fitting a mixture model of normal distributions to K_s_ estimates between homeologs yielded two distinct, but overlapping Gaussian distributions. This suggests two whole genome duplication events are included in our age distribution data. Using a mutation rate calibration for conifer Ks divergence, we estimated the timing of the first whole genome duplication in *Sequoia* to have occurred around 10 Ma, with the second occurring more recently, about 3 Ma. These dates are in apparent contradiction to the discovery of *Sequoia* fossils in the Eocene (33-53 Ma) with guard cells of a size taken to be indicative of polyploidy (Ma et al., 2005). One possible explanation for this discrepancy is that the mutation rate is three-fold lower in *Sequoia* (or redwoods in general) than in other conifers. However, although some redwoods may have extremely long life spans, such a great different in the rate of synonymous substitutions seems improbable.

A second possibility is that the Eocene fossils represent an independent instance of polyploidy in a closely related lineage that was misclassified as being in *Sequoia*. It is noteworthy that some plant groups that acquire the propensity to undergo polyploidy, do so repeatedly, a possible case in point being the *Ephedra* lineage, which appears to have experienced multiple whole genome duplication events (Ickert-Bond, 2003). Further evaluating this hypothesis would require measurements of guard cells in a much larger number of different aged *Sequoia* fossils from different geographic locations.

The final possible explanation for the low divergence of putative homeologs in *Sequoia* is that while autopolyploidy occurred in the Eocene (or even earlier), multisomic inheritance persisted for a long period of time, possibly even to the present for some loci. In such a case the gene duplication events we dated would not correspond to the polyploidy event per se but would reflect subsequent, recombinational homogenization. This hypothesis is consistent with multivalent formation in modern *Sequoia*, and suggests a very slow diploidization process following whole genome duplication in *Sequoia.*

### Implications for polyploidization patterns in gymnosperms

Given what we know about polyploidy in *Sequoia*, what conclusions can we draw about patterns of polyploidization in gymnosperms overall? With the exception of *Ephedra,* instances of polyploid gymnosperms are limited to monospecific genera (e.g. *Sequoia, Fitzroya*), or even just to polyploid individuals within diploid species (e.g. *Juniperus x pftizeriana*; Ahuja, 2005). If polyploidy in gymnosperms is associated with small clades, as seems to be the case, we can infer that polyploidy either hinders speciation or promotes extinction of gymnosperm lineages, or both.

The apparent mismatch between the inferred age of gene duplication and the timing of polyploidization as seen in the fossil record suggests an intriguing hypothesis to explain the paucity of polyploidy in gymnosperms. Perhaps diploidization happens more slowly in gymnosperms (except perhaps *Ephedra*) than in angiosperms. The main long-term benefits of polyploidy (potential sub- and neo-functionalization of genes) require divergence among homeologous chromosomes, which can only happen once loci are diploidized. Thus, continued multisomic inheritance precludes the emergence of any evolutionary advantage in polyploid lineages.

If polyploidy in gymnosperms is more burden than boon, the persistence of hexaploid *Sequoia* may reflect an ability to avoid extinction rather than superior fitness. In this regard it is perhaps noteworthy that *S. sempervirens* manifests some traits that might help stave of extinction, namely clonal reproduction, self-compatibility, and extreme longevity. In coast redwood populations, suckers often emerge from the base of adult trees, extending generation time (meiosis-to-meiosis) almost indefinitely. Furthermore, production of asexual stands may lead to abundant genetic selfing among clonal ramets, as coast redwoods are self-compatible (Burns & Honkala, 1990). This means that a spontaneous polyploid, perhaps gaining the transient advantage of fixed heterozygosity, could spread by a combination of asexual reproduction and selfing. It is conceivable, therefore, that even after the erosion of fixed heterozygosity the lineage could persist despite never gaining the long-term advantages typically associated with polyploidy, instead suffering the concomitant problem of enlarged genome size. The only other natural polyploid in Cupressaceae, *Fitzroya cupressoides,* is a putative autotetraploid. Like *Sequoia, Fitzroya* is both long-lived and capable of clonal reproduction (Silla et al., 2002). Thus, while more work is needed to evaluate the occurrence of multisomic inheritance in both polyploid species (e.g. *Sequoia, Fitzroya*) and polyploid clones *Juniperus xpftizeriana,* our hypothesis can both explain the rarity of neopolyploidy in gymnosperms and why *Sequoia* is an exception to this general rule.

## Acknowledgements

We thank Cécile Ané, Matt Johnson, and Nisa Karimi for improving this manuscript through countless discussions. Support for this project was provided by a grant to ADS and DB from Save The Redwoods League. This material is based upon work supported by the National Science Foundation Graduate Research Fellowship under Grant No. DGE-0718123 to ADS. The data analyses were partly performed using resources provided by the Swedish National Infrastructure for Computing (SNIC) at UPPMAX.

## Author Contributions

ADS and DB designed the research and wrote the manuscript, ADS collected the data, ADS and NS analyzed the data.

